# sORF-Trans2MS: a two-module deep learning framework for sORF translation and MS-supported microprotein prediction

**DOI:** 10.64898/2026.09.29.755517

**Authors:** Jiapeng He, Jingchen Sui, Qiufen Chen, Hailiang Hu, Chris Soon Heng Tan

**Affiliations:** Department of Chemistry, College of Science, Southern University of Science and Technology, Shenzhen, Guangdong, China, 518055; Department of Biochemistry, SUSTech Homeostatic Medicine Institute, School of Medicine, Southern University of Science and Technology, Shenzhen, Guangdong, China, 518055

**Keywords:** Small open reading frames (sORFs), Microproteins, Ribosome profiling, Mass spectrometry, Deep learning

## Abstract

Ribosome profiling is widely used to identify translated small open reading frames (sORFs), but existing prediction models often generalize poorly to newly collected sORF datasets. In addition, many microproteins with strong mass spectrometry (MS) evidence lack support in public Ribo-seq resources, suggesting that Ribo-supported sORF and MS-supported microprotein detection may represent complementary prediction tasks. Here, we developed sORF-Trans2MS, a two-module deep learning framework for predicting Ribo-supported sORFs and MS-supported microproteins. Module I uses RNA-FM representations for upstream, ORF, and downstream regions with a Transformer encoder for Ribo-supported sORF translation prediction. Module II integrates pretrained RNA-FM representations of RNA context with ESM-2 protein representations using multi-scale CNN encoders for predicting translated microproteins supported by MS but not Ribo-seq data. Module I was trained on 9,579 Ribo-supported sORFs and matched background sORFs, while Module II used 14,697 MS-supported (MS^++^Ribo^−^) microproteins and matched backgrounds. Module I achieved an AUROC of 0.870 internally and 0.930 on an independent GENCODE dataset. Ablation analysis highlighted upstream RNA context, including a distinct signal around 50 nt upstream of AUG. Module II achieved an AUROC/AUPR of 0.804/0.797 internally and 0.785/0.773 on an independent mass spectrometry dataset. Model interpretation highlighted upstream RNA context and the protein N terminus, while literature-supported microproteins further illustrated the complementary roles of the two modules. Overall, sORF-Trans2MS improves prediction and generalization of Ribo-supported sORFs and extends computational discovery to MS-supported candidates that are not represented in current public Ribo-seq resources.

## Introduction

Small open reading frames (sORFs) and their encoded microproteins represent an important but still incompletely annotated part of the human proteome. Microproteins are small proteins that are encoded by short open reading frames (sORFs) located outside conventional protein coding regions, often no more than 150 amino acids in length (Leong et al. 2022; Jaunbocus et al. 2025). Because of their small size, weak sequence conservation, and low abundance, many sORFs have been overlooked by traditional genome annotation pipelines (Delcourt et al. 2018). Increasing evidence has shown that microproteins play important role in diverse biological processes, including metabolism, development, stress responses, and pathways related to cancer (Leong et al. 2022). For example, DWORF and myoregulin regulate SERCA-dependent calcium handling and muscle function (Anderson et al. 2015; Nelson et al. 2016), SPAR modulates mTORC1 signaling and muscle regeneration (Matsumoto et al. 2017), NoBody interacts with the mRNA decapping complex and regulates cytoplasmic P-body abundance (D’Lima et al. 2017). These findings suggest that microproteins are a functionally important but still underexplored component of the proteome.

Ribosome profiling has greatly improved the discovery of translated sORFs by sequencing mRNA fragments protected by ribosomes at near nucleotide resolution (Ingolia et al. 2009). Tools such as ribotricer further use the triple nucleotide periodicity of Ribo-seq reads to identify actively translated ORFs, including short ORFs (Choudhary et al. 2020). However, the detection of sORFs from Ribo-seq remains sensitive to data quality, sequencing depth, and sparse ribosome footprint coverage (Chothani et al. 2023). Mass spectrometry provides complementary evidence at the protein level, but many noncanonical proteins remain difficult to detect because of their short length, low abundance, and limited peptide detectability (Wacholder and Carvunis 2023). Together, Ribo-seq and mass spectrometry provide complementary evidence for microprotein discovery, but neither method alone captures the complete microproteome.

Several databases and computational tools have been developed to support sORF annotation. OpenProt provides expanded protein annotations, including reference proteins, alternative proteins, and isoforms with supporting evidence from translation and proteomics data (Brunet et al. 2019; Leblanc et al. 2024). SmProt collects microproteins from ribosome profiling, literature, and other evidence sources (Li et al. 2021). In addition, a number of computational methods have been developed to predict coding potential from sequence features and have also been applied to sORF prediction. For example, MiPepid is a machine learning tool designed to identify sORFs from nucleotide sequences (Zhu and Gribskov 2019). DeepCPP is a deep learning model for predicting RNA coding potential using nucleotide bias information (Zhang et al. 2021), while sOCP predicts coding potential of sORFs using translation initiation site and in-frame features (Peng et al. 2024b). However, with the rapid increase in newly reported microproteins and sORF candidates in recent years, these models showed limited generalization on newly curated sORF candidates. Their limited performance motivated us to develop a model for distinguishing Ribo-seq supported sORFs from background candidates.

Another challenge is that current microprotein studies largely focus on candidates supported by Ribo-seq, particularly those with additional mass spectrometry evidence. Using a microprotein enrichment method developed in our laboratory (Li et al. 2026), we identified a group of microproteins supported by strong mass spectrometry evidence (stringent FDR of 0.01% and PSM filtering with PepQuery) but without corresponding Ribo-seq support in public databases (Li et al. 2026). This observation suggests that current Ribo-seq datasets may not capture all translated sORFs and that some microproteins can be missed when discovery relies mainly on Ribo-seq evidence. To address this gap, we developed a model specifically for MS^++^Ribo^−^ microproteins and evaluated its performance on an independent mass spectrometry dataset.

Recent advances in biological language models provide useful representations for both RNA and protein sequences. RNA Foundation Model (RNA-FM) is a self-supervised RNA foundation model trained on large-scale RNA sequences and can provide contextual RNA sequence representations related to RNA structure and function (Chen et al. 2022). Its utility for RNA structural modeling has also been demonstrated in RhoFold+ (Shen et al. 2024). RNA structure and sequence context in the 5′untranslated region are known to influence translational regulation (Razumova et al. 2025). We therefore used RNA-FM representations followed by a Transformer encoder to capture information across an extended RNA context. On the protein side, ESM-2 learns residue representations from large-scale protein sequence data and captures information relevant to protein structure (Lin et al. 2023). These representations provide complementary RNA and protein features for microprotein prediction.

Here, we present sORF-Trans2MS, a two-module deep learning framework for sORF translation and MS-supported microprotein prediction. Rather than treating all experimentally supported sORFs as a single positive class, we considered Ribo-seq supported translation and MS-supported microprotein detection as two complementary prediction tasks. In Module I, RNA-FM is used to obtain contextual token representations from the upstream, ORF, and downstream regions, providing sequence features associated with RNA structure and function. The resulting representations are combined with relative position information and further refined by a Transformer encoder. This design enables long-range sequence modeling, which is particularly suitable for RNA because folding can create structural relationships between positions that are distant in the linear sequence. The refined features are then pooled and passed to an MLP classifier for Ribo-supported sORF translation prediction. Module II integrates RNA-FM representations of the three RNA regions with ESM-2 residue representations of the encoded protein. RNA and protein features are processed by separate position-aware CNN encoders. The refined RNA and protein features are subsequently fused and passed to an MLP classifier for MS^++^Ribo^−^ microprotein prediction. Together, the two modules provide complementary modeling strategies for Ribo-supported translation and MS-supported microprotein detection, including candidates that may not be captured by current public Ribo-seq resources.

## Results

We developed sORF-Trans2MS as a two-module framework for Ribo-supported sORF translation and MS-supported microprotein prediction (**Figure 1A**). Module Ⅰ uses RNA context to predict Ribo-supported translation (**Figure 1B**), whereas Module Ⅱ integrates RNA and protein sequence information to predict MS-supported microproteins lacking public Ribo-seq support (**Figure 1C**). The two modules were trained independently and generated separate prediction scores.

**Figure 1.**
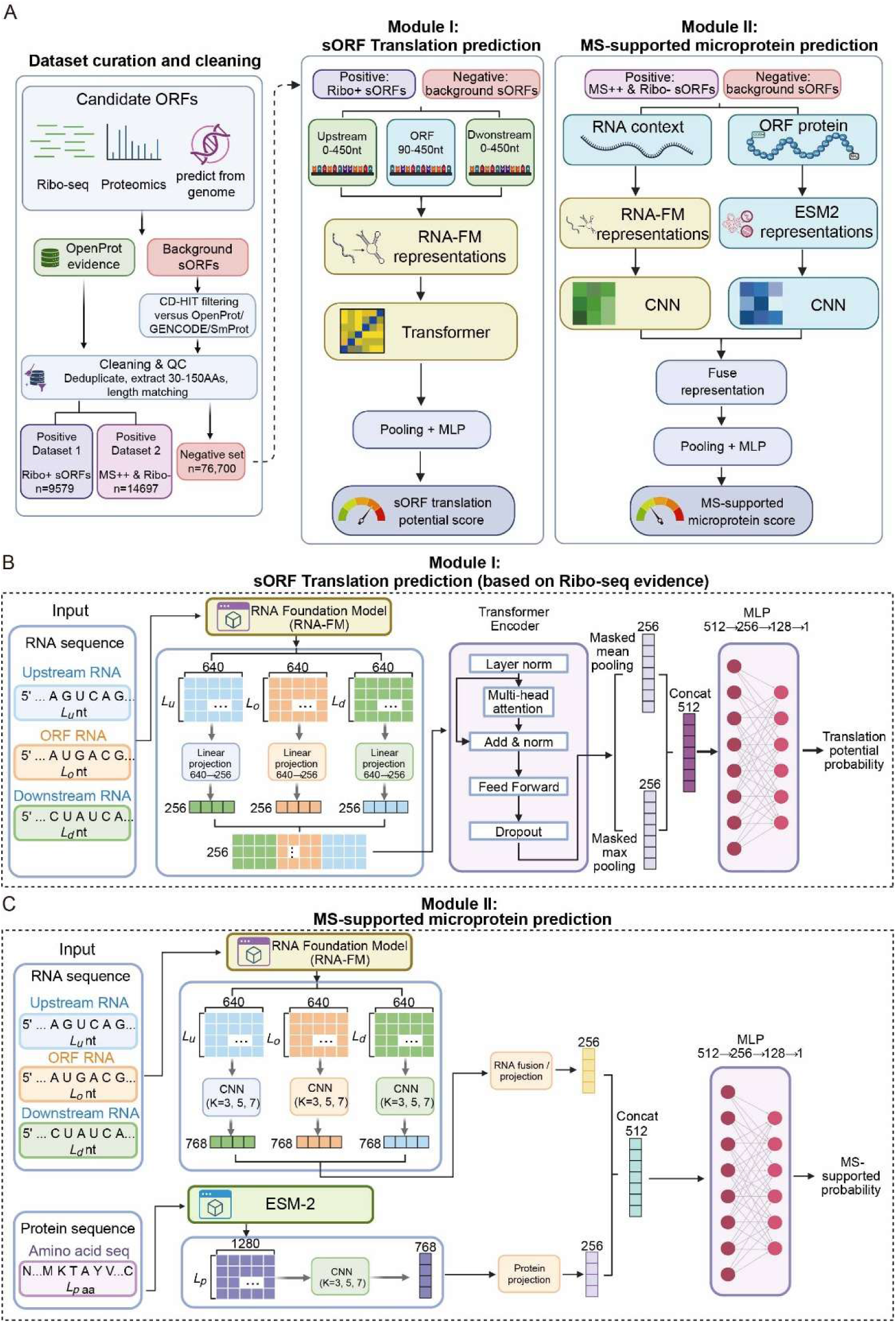
Overview of dataset construction and the sORF-Trans2MS framework. **A**. Construction of Ribo+ and MS++Ribo− positive datasets and the background sORF dataset. Candidate background ORFs were filtered against OpenProt, GENCODE, and SmProt, followed by quality control, deduplication, length filtering, and length matching. **B**. Architecture of Module I for Ribo-supported sORF translation prediction. RNA-FM representations of the upstream, ORF, and downstream regions are combined with relative position information and processed by a Transformer encoder, followed by sequence pooling and an MLP classifier. **C**. Architecture of Module II for MS-supported microprotein prediction. RNA-FM and ESM-2 representations are processed by separate multi-scale CNN encoders, projected into RNA and protein representations, concatenated, and passed to an MLP classifier.

### Rationale for separating Ribo-supported and MS-supported sORF prediction

To examine whether Ribo-seq-supported and MS-supported sORFs could be modeled within a single prediction task, we first grouped candidates according to the Ribo-seq and mass spectrometry (MS) evidence annotations provided by OpenProt v2.2. In OpenProt, MS scores of 0, 1 and ≥2 were labeled as MS^−^, MS^+^ and MS^++^, respectively, whereas Translation Event (TE) scores of 0, 1 and ≥2 were labeled as Ribo^−^, Ribo^+^ and Ribo^++^, respectively. Compared with the Ribo-supported dataset, adding MS^++^Ribo^−^ candidates to the Ribo-supported positive set substantially reduced performance, with AUROC decreasing from 0.870 to 0.763 and AUPR from 0.874 to 0.762 (**Figure 2A, B**). These results indicate that Ribo-supported and MS-supported candidates do not form a homogeneous positive class.

**Figure 2.**
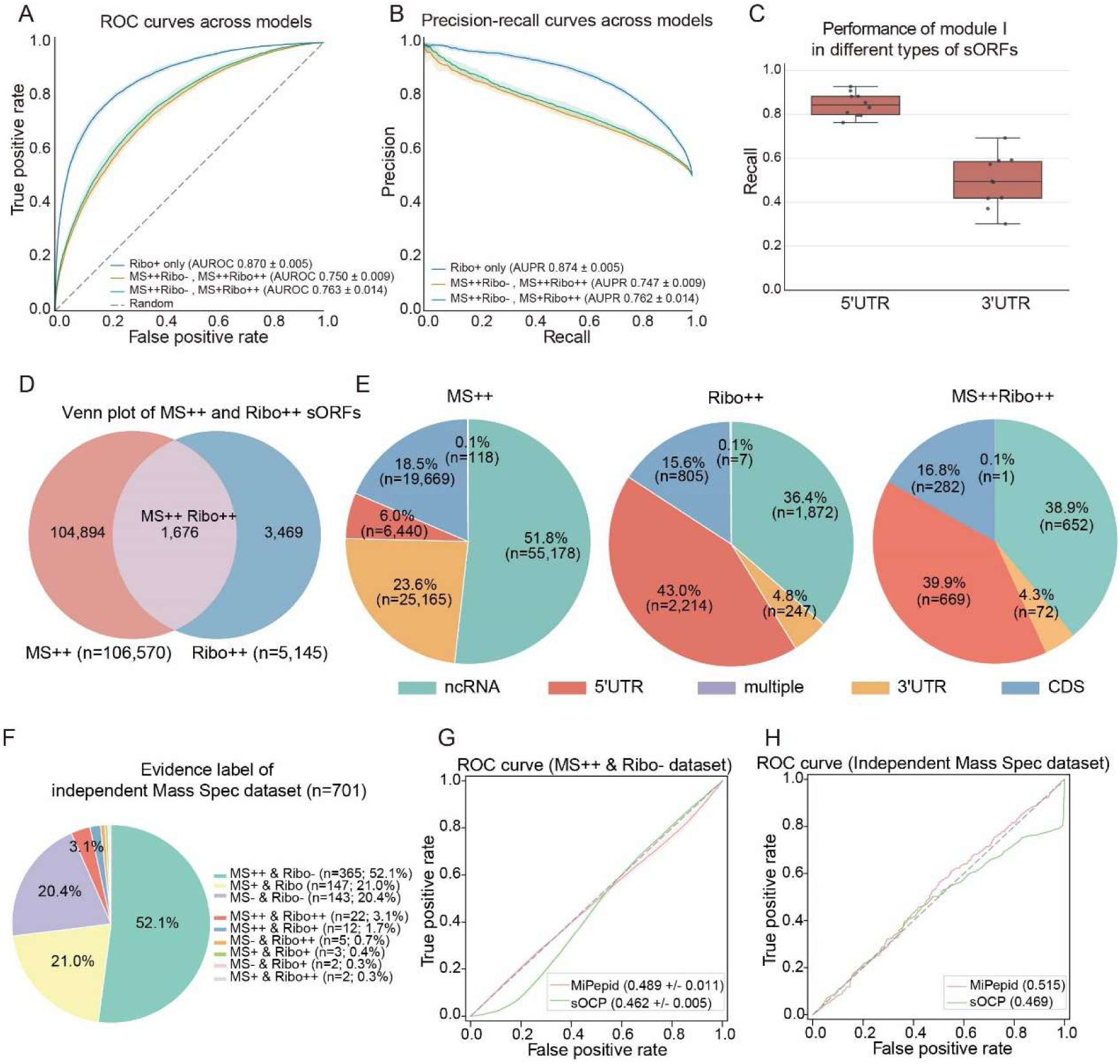
Rationale for separating Ribo-supported and MS-supported sORF prediction. **A, B**. ROC and PR curves comparing models trained on the Ribo-supported (Ribo^++^) dataset and mixed datasets containing MS^++^Ribo^−^ candidates. **C**. Recall of Module I for 5′UTR- and 3′UTR-derived sORFs. **D**. Overlap between MS^++^ and Ribo^++^ sORFs. **E**. Transcript-region distributions of MS^++^, Ribo^++^, and MS^++^Ribo^++^ sORFs. **F**. Distribution of Ribo-seq and mass spectrometry evidence in the independent mass spectrometry dataset. **G, H**. ROC curves of MiPepid, and sOCP on the MS^++^Ribo^−^ dataset and the independent mass spectrometry dataset, respectively.

We then examined whether this difference was associated with transcriptomic location. Module I showed substantially higher recall for 5′UTR-derived than for 3′UTR-derived sORFs (**Figure 2C**). Consistently, MS^++^ and Ribo^++^ candidates showed markedly different regional distributions (**Figure 2D, E**). MS^++^ sORFs contained a higher proportion of 3′UTR candidates (MS^++^: 23.6% vs Ribo^++^: 4.8%), whereas Ribo^++^ sORFs were more strongly enriched in 5′UTR (MS^++^: 6.0% vs Ribo^++^: 43.0%) (**Figure 2E**). Candidates supported by both MS^++^ and Ribo^++^ evidence showed a distribution more similar to the Ribo^++^ group.

This distinction was further supported by an independent MS-supported microprotein dataset assembled from the two published proteomic studies (Li et al. 2022a; Li et al. 2026), in which 655 of 701 microproteins lacked public Ribo-seq support (**Figure 2F**). Published sORF prediction tools showed limited performance on internal MS^++^Ribo^-^ dataset and independent MS dataset. (**Figure 2G,H**). Together, these results indicate that Ribo-supported translation and MS-supported microprotein detection represent related but distinct prediction tasks. We therefore modeled these two tasks separately as Module I and Module II.

### Performance evaluation of Module I for sORF translation prediction

Module I combines RNA-FM representations of upstream, ORF and downstream regions with a Transformer encoder and MLP classifier. It was trained to distinguish Ribo-supported sORFs from background candidates. We then compared Module I with baseline models and existing sORF prediction tools on the internal test set (**Table 1**; **Figure 3A, B**). Across ten independent runs, Module I achieved an AUROC of 0.870 ± 0.005 and an AUPR of 0.874 ± 0.005. The CNN model showed comparable performance, with slightly higher AUROC and AUPR values of 0.875 ± 0.006 and 0.876 ± 0.007, respectively. The other baseline models generally showed lower performance. Among the published sORF prediction tools, MiPepid and sOCP achieved AUROCs of 0.629 and 0.489, respectively, indicating limited performance on the newly constructed benchmark.

**Figure 3.**
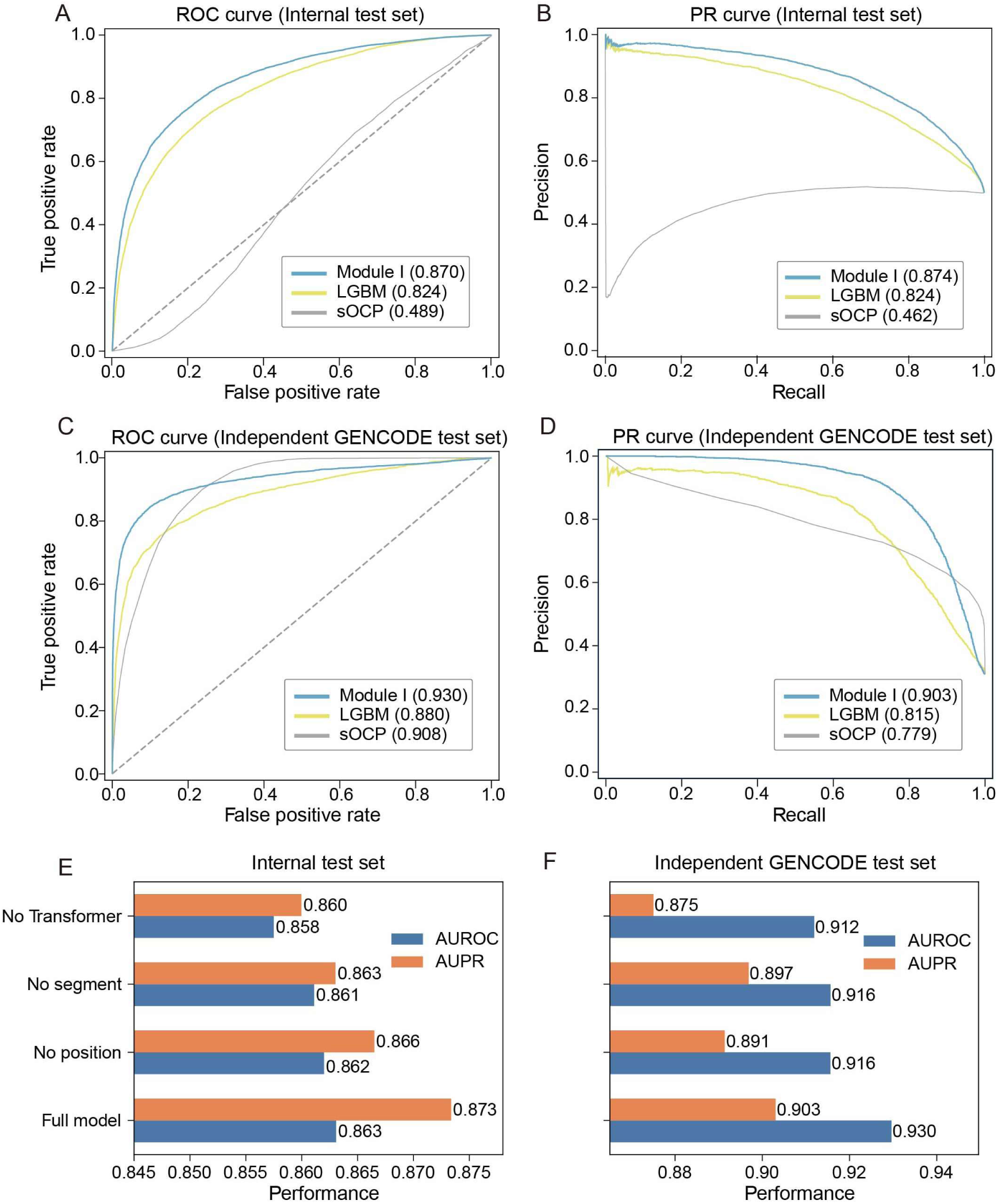
Performance of Module I for sORF translation prediction. **A**. ROC curves of different models on the internal test set. Dashed lines in the ROC plots indicate random prediction. **B**. PR curves of different models on the internal test set. **C**. ROC curves of different models on the independent GENCODE test set. Dashed lines in the ROC plots indicate random prediction. **D**. PR curves of different models on the independent GENCODE test set. **E**, **F.** Architecture ablation analysis of Module I on the internal test set and independent GENCODE test set. Bar plot displays performance of the full model and variants without positional features, segment embeddings or the Transformer encoder.

**Table 1.** Performance Comparison of Different Models for Module I on the Internal Test Set.

| dataset | model | Accuracy | Precision | Recall | F1 | AUROC | AUPR |
| --- | --- | --- | --- | --- | --- | --- | --- |
| Internal_test | **Module I** | **$0.782 \pm$** | **$0.834 \pm$** | $0.710 \pm$ | $0.763 \pm$ | $0.870 \pm$ | $0.874 \pm$ |
|  |  | **0.017** | **0.032** | 0.078 | 0.035 | 0.005 | 0.005 |
| Internal_test | LGBM | $0.748 \pm$ | $0.773 \pm$ | $0.702 \pm$ | $0.735 \pm$ | $0.824 \pm$ | $0.824 \pm$ |
|  |  | 0.009 | 0.012 | 0.013 | 0.010 | 0.008 | 0.012 |
| Internal_test | CNN | 0.786 ± | 0.818 ± | 0.747 ± | **0.776 ±** | **0.875 ±** | **0.876 ±** |
|  |  | 0.013 | 0.046 | 0.087 | **0.030** | **0.006** | **0.007** |
| Internal_test | Logistic | 0.671 ± | 0.673 ± | 0.668 ± | 0.670 ± | 0.733 ± | 0.722 ± |
|  | Regression | 0.010 | 0.014 | 0.008 | 0.007 | 0.013 | 0.017 |
| Internal_test | Random Forest | 0.731 ± | 0.782 ± | 0.642 ± | 0.705 ± | 0.806 ± | 0.808 ± |
|  |  | 0.006 | 0.013 | 0.014 | 0.007 | 0.005 | 0.007 |
| Internal_test | pooled MLP | 0.753 ± | 0.761 ± | 0.737 ± | 0.749 ± | 0.833 ± | 0.831 ± |
|  |  | 0.004 | 0.007 | 0.012 | 0.005 | 0.005 | 0.008 |
| Internal_test | MiPepid | 0.511 | 0.505 | **0.98** | 0.666 | 0.629 | 0.614 |
| Internal_test | sOCP | 0.49 | 0.486 | 0.389 | 0.432 | 0.489 | 0.462 |

We next evaluated the models on an independent GENCODE dataset (Deutsch et al. 2026) to examine their performance on newly collected sORFs (**Table 2**; **Figure 3C, D**). Module I achieved an AUROC of 0.930 and an AUPR of 0.903, representing the highest values among the evaluated models. In contrast, the published tools showed less consistent performance. Although sOCP reached an AUROC of 0.908 and a recall of 0.998, its AUPR was lower at 0.779. MiPepid showed lower overall performance on the GENCODE dataset (**Table 2**).

**Table 2.** Performance Comparison of Different Models for Module I on the Independent GENCODE Test Set.

| dataset | model | Accuracy | Precision | Recall | F1 | AUROC | AUPR |
| --- | --- | --- | --- | --- | --- | --- | --- |
| Gencode | **Module I** | **0.875** | **0.766** | 0.857 | **0.809** | **0.93** | **0.903** |
| external test |  |  |  |  |  |  |  |
| Gencode | LGBM | 0.814 | 0.667 | 0.794 | 0.725 | 0.88 | 0.815 |
| external test |  |  |  |  |  |  |  |
| Gencode | CNN | 0.836 | 0.701 | 0.821 | 0.756 | 0.908 | 0.842 |
| external test |  |  |  |  |  |  |  |
| Gencode | Logistic | 0.642 | 0.435 | 0.517 | 0.472 | 0.643 | 0.468 |
| external test | Regression |  |  |  |  |  |  |
| Gencode | Random Forest | 0.847 | 0.729 | 0.807 | 0.766 | 0.91 | 0.818 |
| external test |  |  |  |  |  |  |  |
| Gencode | pooled MLP | 0.773 | 0.611 | 0.743 | 0.67 | 0.846 | 0.76 |
| external test |  |  |  |  |  |  |  |
| Gencode | MiPepid | 0.309 | 0.292 | 0.859 | 0.436 | 0.578 | 0.435 |
| external test |  |  |  |  |  |  |  |
| Gencode | sOCP | 0.595 | 0.432 | **0.998** | 0.603 | 0.908 | 0.779 |
| external test |  |  |  |  |  |  |  |

Architecture ablation further showed reduced performance after removal of positional features, segment embeddings or the Transformer encoder (**Figure 3E, F**).

### Ablation analysis reveals important upstream RNA context in Module I

To identify which RNA regions contributed most to Module I prediction, we performed region-level ablation of the upstream, ORF, and downstream sequences. Masking the upstream region caused the largest decrease in both AUROC and AUPR across ten runs, indicating that the upstream sequence provided the strongest contribution to translation prediction (**Figure 4A, C**). We next performed position-level ablation across the upstream region to further localize the informative signals (**Figure 4B, D**). Positions close to the AUG start codon showed a clear contribution, consistent with the known importance of the translation initiation context (Hernandez et al. 2024). Notably, a broader signal was also observed approximately −60 to - 25nt upstream of the AUG, with an importance comparable in magnitude to the AUG-proximal signal. This pattern was consistently observed using both AUROC and AUPR as evaluation metrics (**Figure 4B, D**). To investigate the sequence features underlying this upstream signal, we performed de novo motif analysis across representative upstream windows. De novo motif analysis revealed recurrent C/G-rich sequence patterns across multiple upstream regions (**Figure 4E**). Notably, the −60 to −25 nt window contained a strongly enriched CCGCCGCSG motif (E = 6.4 × 10⁻¹⁵), which contains the previously reported CCGCCGCCG sequence associated with eIF4A-dependent translation (**Figure 4E, Table S1**) (Singh et al. 2021). In addition, CUGMG motif wasspecifically enriched within the −60 to −25 nt window (E = 4.6×10^-2^) and showed consistent enrichment across 10 independent runs (**Figure 4F**). We additionally compared predicted RNA secondary-structure features between positive and negative sORFs. Only modest differences in paired fraction and structural-class frequencies were observed across the upstream region (**Figure S1**).

**Figure 4.**
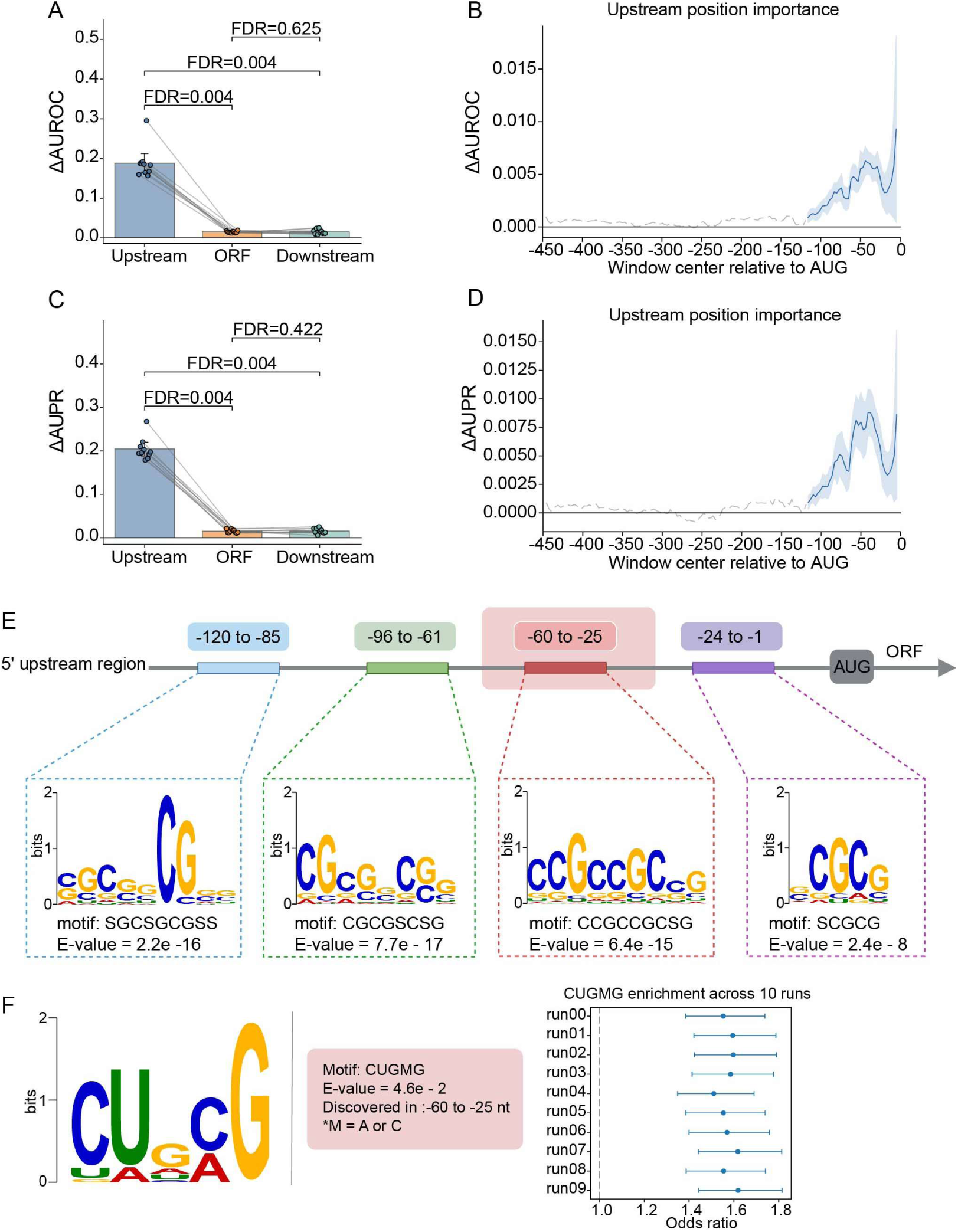
Ablation analysis of RNA sequence context in Module I. **A**. Region level ablation of upstream, ORF, and downstream RNA sequences, evaluated by decreases in AUROC across ten runs. **B**. Position level ablation of the upstream region, showing changes in AUROC relative to the AUG start codon. The region with bold line indicates positions covered by at least 80% of the evaluated sequences. **C**. Region level ablation of upstream, ORF, and downstream RNA sequences, evaluated by decreases in AUPR across ten runs. **D**. Position level ablation of the upstream region, showing changes in AUPR relative to the AUG start codon. The blue line indicates positions covered by at least 80% of the evaluated sequences. Positive ΔAUROC or ΔAUPR values indicate decreased model performance after ablation and therefore greater feature importance. Statistical comparisons were performed using paired two-sided Wilcoxon rank tests with Benjamini Hochberg FDR correction. **E**. De novo motif analysis of four upstream regions. The most significantly enriched motif identified by STREME is shown for each window. **F**. Identification and validation of motif specifically enriched in −60 to −25nt upstream region. The sequence logo shows the CUGMG motif identified by STREME in the discovery run (E = 4.6 × 10⁻²; M = A or C). The forest plot shows enrichment of CUGMG motif in each run.

### Performance evaluation of Module II for MS-supported microprotein prediction

Module II integrates RNA-FM representations of the upstream, ORF and downstream regions with ESM-2 protein representations. The RNA and protein features are processed by separate CNN encoders and fused for MS-supported microprotein prediction. We then evaluated Module II for MS^++^Ribo^−^ microprotein prediction. On the internal test set, Module II achieved an AUROC of 0.804 ± 0.006 and an AUPR of 0.797 ± 0.007 across ten independent runs (**Table 3**; **Figure 5A**). The Transformer achieved AUROC of 0.764 ± 0.012 and AUPR of 0.728 ± 0.015. Classical models, including pooled MLP, LR, RF, and LightGBM, generally showed lower performance (**Table 3**). Published sORF prediction tools performed close to random, with AUROC of 0.489 for MiPepid and 0.462 for sOCP. Overall, Module II showed better performance than the classical baselines and published sORF prediction tools on the internal test set.

**Table 3.** Performance Comparison of Different Models for Module II on the Internal Test Set.

| dataset | model | Accuracy | Precision | Recall | F1 | AUROC | AUPR |
| --- | --- | --- | --- | --- | --- | --- | --- |
| Module2_inte | **Module II** | **0.716 ±** | **0.748 ±** | 0.660 ± | **0.697 ±** | **0.804 ±** | **0.797 ±** |
| rnal_test |  | **0.013** | **0.029** | 0.086 | **0.037** | **0.006** | **0.007** |
| Module2_inte | Transformer | 0.695 ± | 0.696 ± | 0.696 ± | 0.695 ± | 0.764 ± | 0.728 ± |
| rnal_test |  | 0.012 | 0.021 | 0.036 | 0.013 | 0.012 | 0.015 |
| Module2_inte | pooled MLP | 0.662 ± | 0.643 ± | **0.735 ±** | 0.684 ± | 0.731 ± | 0.706 ± |
| rnal_test |  | 0.006 | 0.018 | **0.065** | 0.019 | 0.006 | 0.007 |
| Module2_inte | Logistic | 0.607 ± | 0.607 ± | 0.611 ± | 0.609 ± | 0.650 ± | 0.620 ± |
| rnal_test | Regression | 0.006 | 0.006 | 0.008 | 0.006 | 0.007 | 0.007 |
| Module2_inte | Random Forest | 0.652 ± | 0.636 ± | 0.711 ± | 0.671 ± | 0.710 ± | 0.684 ± |
| rnal_test |  | 0.006 | 0.005 | 0.010 | 0.006 | 0.008 | 0.009 |
| Module2_inte | LGBM | 0.657 ± | 0.642 ± | 0.709 ± | 0.674 ± | 0.721 ± | 0.698 ± |
| rnal_test |  | 0.006 | 0.006 | 0.012 | 0.006 | 0.006 | 0.009 |
| Module2_inte | Module I | 0.571 ± | 0.672 ± | 0.288 ± | 0.395 ± | 0.622 ± | 0.626 ± |
| rnal_test |  | 0.018 | 0.041 | 0.087 | 0.082 | 0.016 | 0.016 |
| Module2_inte | MiPepid | 0.490 ± | 0.395 ± | 0.731 ± | 0.512 ± | 0.489 ± | 0.497 ± |
| rnal_test |  | 0.006 | 0.208 | 0.385 | 0.270 | 0.011 | 0.010 |
| Module2_inte | sOCP | 0.461 ± | 0.443 ± | 0.297 ± | 0.356 ± | 0.462 ± | 0.447 ± |
| rnal_test |  | 0.005 | 0.008 | 0.008 | 0.008 | 0.005 | 0.003 |

We further evaluated Module II on the independent MS dataset (**Table 4**; **Figure 5B**). Module II maintained an AUROC of 0.785 and an AUPR of 0.773. Among the other models, RF also showed relatively strong performance with an AUROC of 0.780, whereas the Transformer, pooled MLP, LR, and LightGBM showed lower overall performance. MiPepid and sOCP performed poorly again, with AUROCs of 0.515 and 0.469, respectively.

**Table 4.**
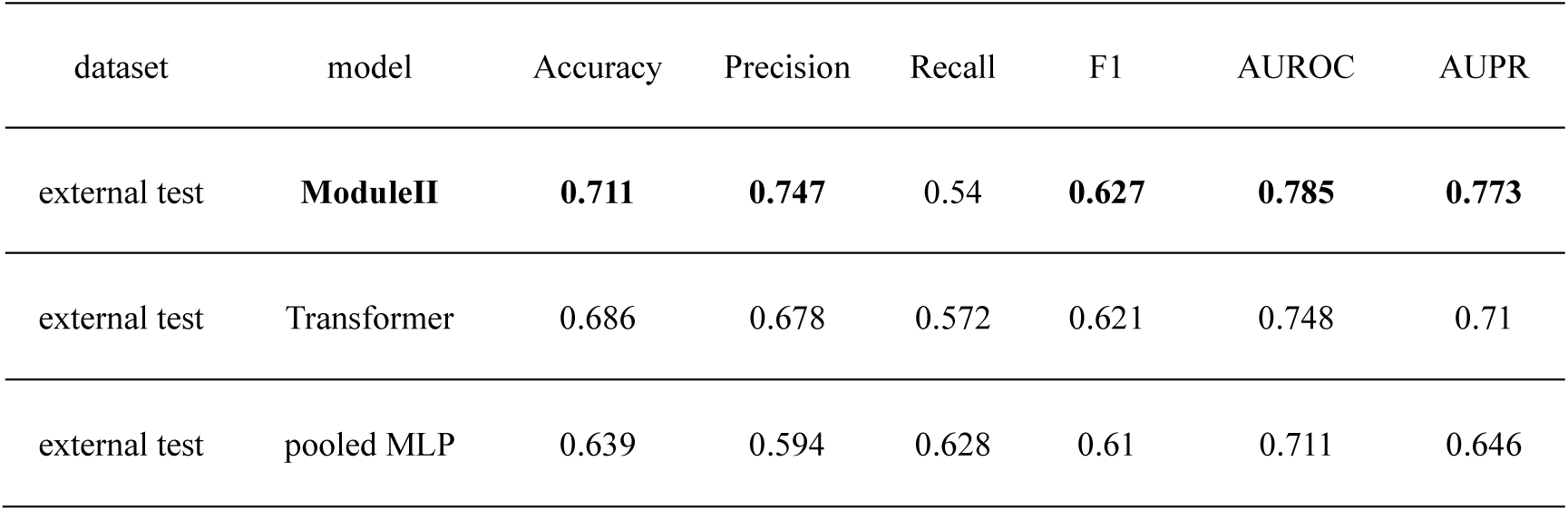
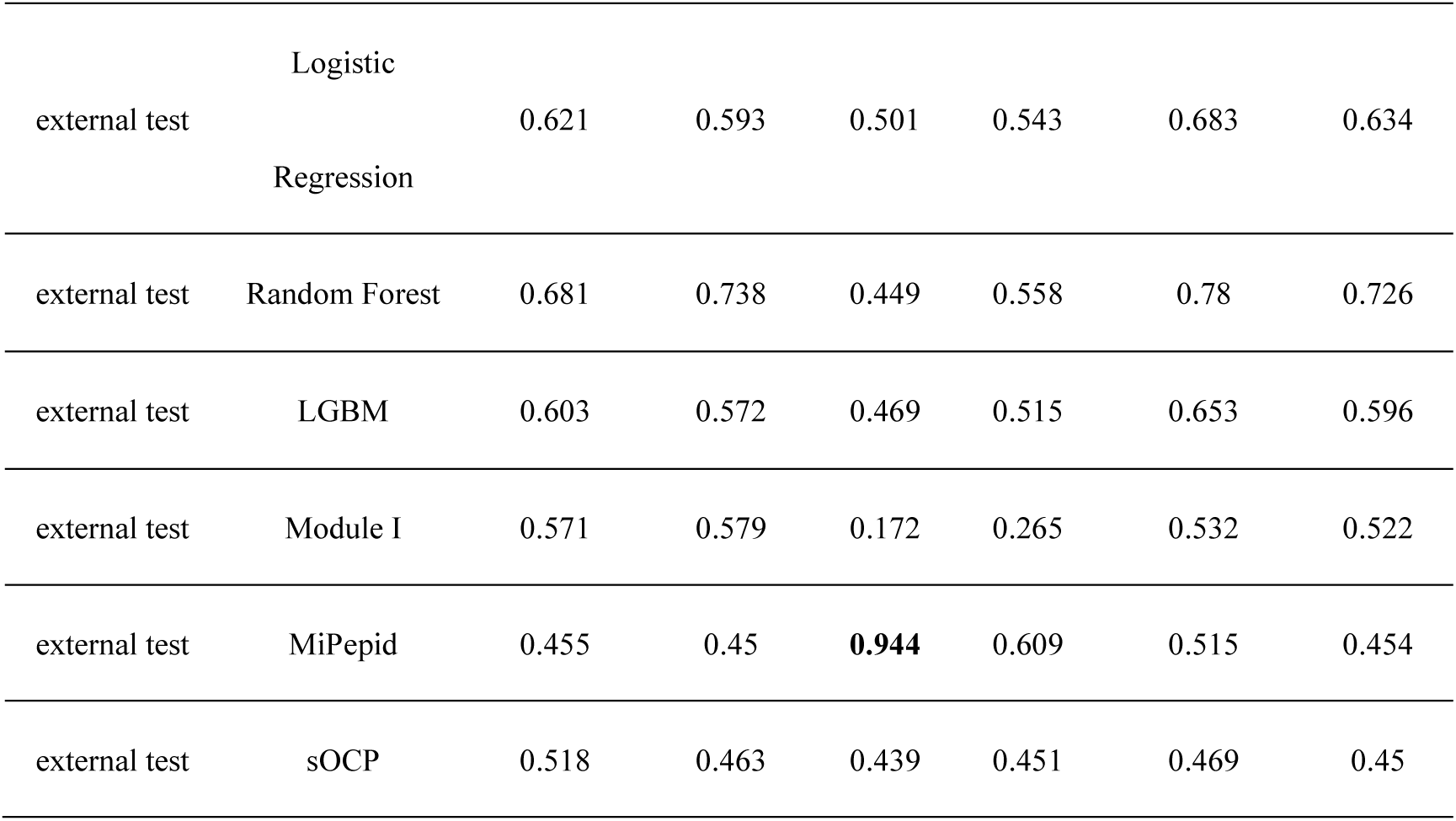
Performance Comparison of Different Models for Module II on the Independent Mass Spectrometry Test Set.

**Figure 5.**
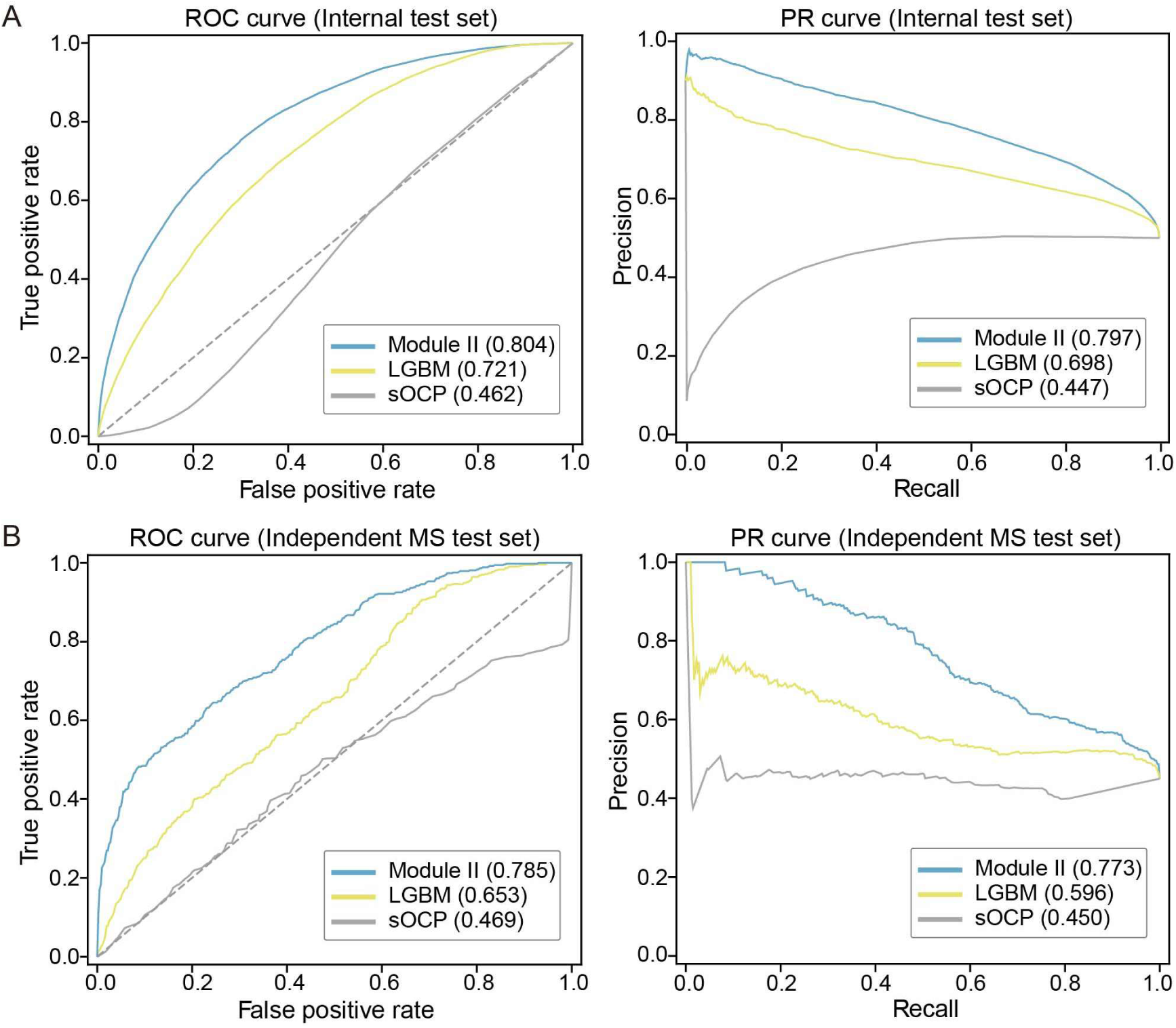
Performance of Module II for MS-supported microprotein prediction. **A**. Receiver operating characteristic (ROC) and precision-recall (PR) curves comparing Module Ⅱ with Transformer, pooled MLP, logistic regression (LR), random forest (RF), LightGBM (LGBM), MiPepid, and sOCP on the internal test set. **B**. ROC and PR curves of the same models on the independent mass spectrometry dataset.

### Ablation analysis identifies key RNA and protein features in Module II

We next examined the contribution of RNA and protein features to Module II prediction. Region-level ablation showed that the upstream RNA and protein features were the major contributors, with protein ablation causing a larger decrease in model performance (**Figure 6A**). Position-level ablation of the upstream RNA region revealed the strongest signal near the AUG start codon, together with a second peak around 50 nt upstream (**Figure 6B; Figure S2B**).

**Figure 6.**
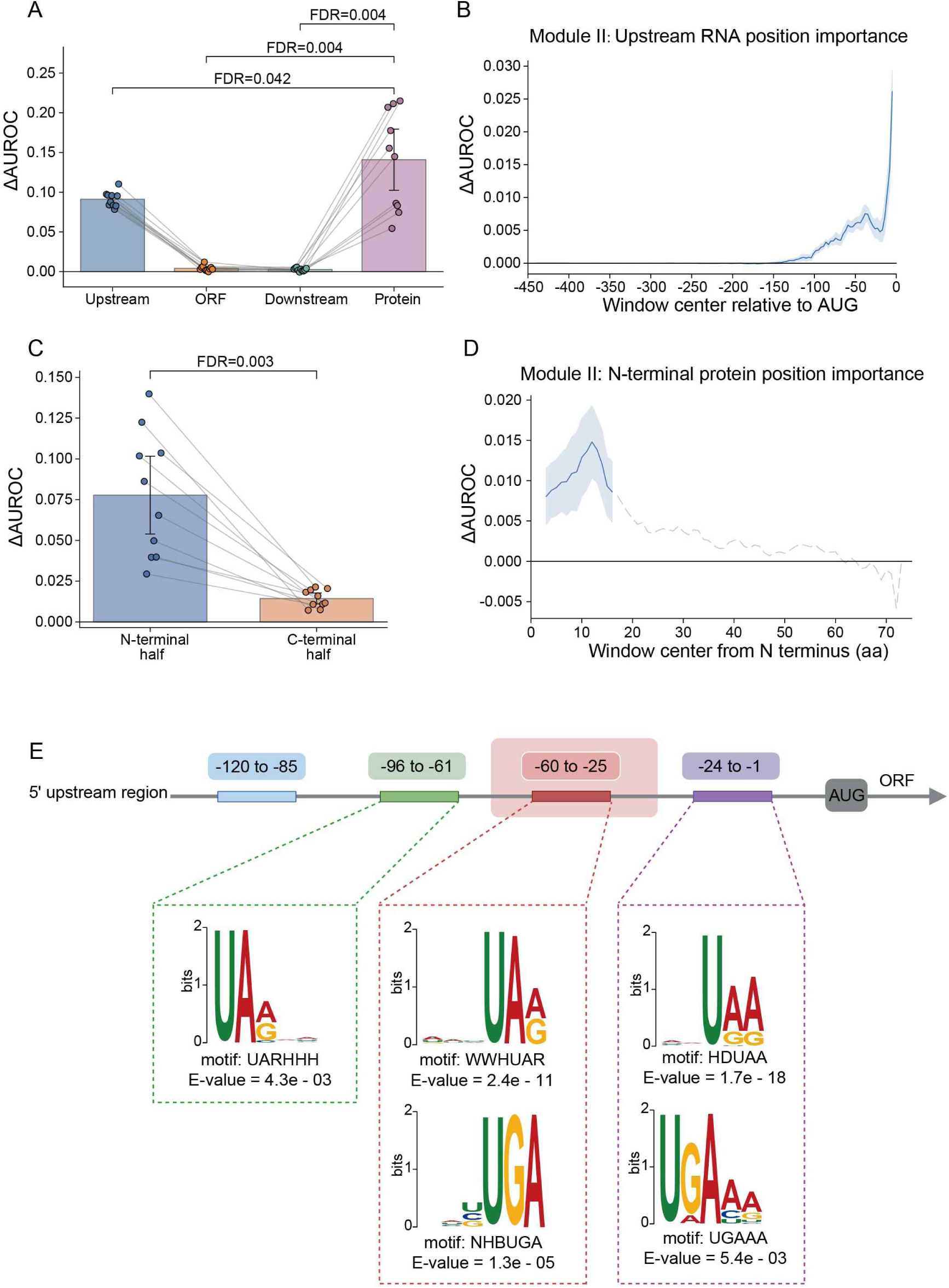
RNA and protein ablation analysis of Module II. **A**. Comparison of ΔAUROC after ablation of the upstream, ORF, downstream, and protein inputs. **B**. Position-level ΔAUROC across the upstream RNA region. **C**. Comparison of ΔAUROC after ablation of the N-terminal and C-terminal halves of the protein sequence. **D**. Position-level ΔAUROC across the N-terminal half of the protein sequence. All of the results are summarized across ten runs. Positive ΔAUROC values indicate decreased model performance after ablation and therefore greater predictive importance. Statistical comparisons were performed using paired two-sided Wilcoxon signed-rank tests with Benjamini-Hochberg correction. **E**. De novo motif analysis of four upstream regions. All significantly enriched motifs identified by STREME are shown for each window.

Unlike Module I, in which the distal upstream signal was comparable to the AUG-proximal signal, the corresponding signal was substantially weaker in Module II. De novo motif analysis further identified distinct sequence patterns across these upstream regions (**Figure 6E, Table S2**). In contrast to the recurrent C/G-rich motifs observed in Module I, Module II predominantly showed A/U-rich motifs, including UARHHH in the −96 to −61 nt region, WWHUAR and NHBUGA in the −60 to −25 nt region, and HDUAA and UGAAA close to the start codon. These results suggest that the two modules capture different upstream sequence features, with Module II showing a stronger dependence on the sequence context close to the AUG.

We then examined which part of the protein sequence contributed most to prediction. Dividing the protein into N-terminal and C-terminal halves showed that ablation of the N-terminal half caused a significantly larger performance decrease than ablation of the C-terminal half (**Figure 6C**). Position-level analysis further localized the strongest contribution to approximately the first 15 to 20 amino acids of the protein (**Figure 6D**). The same overall patterns were observed when feature importance was evaluated using AUPR (**Figure S2**).

### Application of sORF-Trans2MS to literature-supported microproteins

To further evaluate the applicability of sORF-Trans2MS, we collected literature-reported microproteins with experimental Ribo-seq or mass spectrometry evidence (**Figure 7A**). Reported ORF sequences were matched to representative Ensembl transcripts, and the corresponding RNA context and protein sequences were reconstructed using the same preprocessing strategy as the benchmark datasets. Each candidate was then scored independently by Module I and Module II.

**Figure 7.**
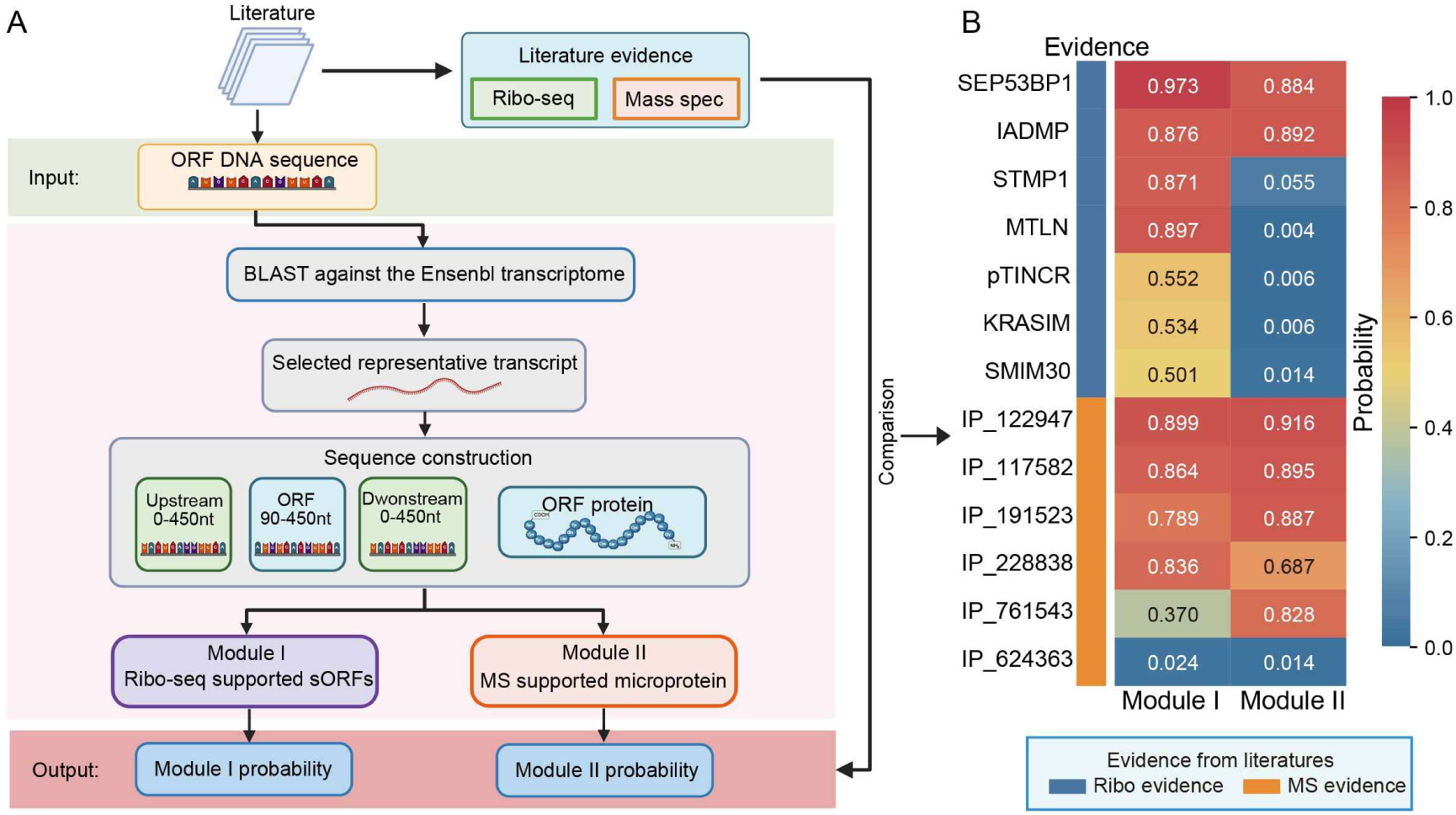
Evaluation of literature-supported sORFs and microproteins using sORF-Trans2MS. **A**. Workflow for reconstructing sequence inputs and evaluating literature-reported candidates with Module I and Module II. Reported ORF sequences were matched to Ensembl transcripts, followed by construction of upstream, ORF, downstream, and protein sequence inputs. **B**. Prediction probabilities of Module I and Module II for candidates supported by Ribo-seq or mass spectrometry evidence in the literature.

All seven candidates with reported Ribo-seq evidence received Module I probabilities of at least 0.5 (**Figure 7B**) (Li et al. 2020; Xu et al. 2020; Li et al. 2022b; Xie et al. 2022; Inchingolo et al. 2023; Yang et al. 2023; Wu et al. 2026). Among six candidates supported by mass spectrometry evidence, five received Module II probabilities above 0.5 (Gagnon et al. 2021; Leblanc et al. 2023; Zhang et al. 2023; Comtois et al. 2025). Several candidates showed high probabilities from both modules, whereas IP_761543 showed a substantially higher Module II probability (0.828) than Module I probability (0.370) (Peng et al. 2024a). These results further illustrate the complementary predictive characteristics of the two modules for experimentally supported microproteins.

## Discussion

Ribosome profiling and mass spectrometry have expanded the catalog of translated sORFs or microproteins. However, existing prediction tools generalized poorly to our newly curated datasets and struggled to identify MS-supported candidates without public Ribo-seq evidence. Our previous proteomic work provided strong evidence that some of these microproteins are expressed (Li et al. 2026), showing that microproteins can be translated even without detectable Ribo-seq support. Moreover, prediction performance decreased when MS-supported candidates and Ribo-supported microproteins were combined into a single positive set. Besides, these two groups differed in their distribution across transcript regions. These findings support the two-module design of sORF-Trans2MS.

Module I showed strong performance on the independent GENCODE dataset relative to published models and internal baselines. This result supports the generalization of the RNA features learned by Module I beyond the training dataset. Ablation analysis further identified an important upstream signal: approximately −60 to −25 nt from the AUG, showing that informative sequence features extend well beyond the start codon. C/G-rich motifs were enriched across multiple upstream regions. They are associated with eIF4A-dependent translation (Waldron et al. 2019; Singh et al. 2021). The CUGMG motif was specifically enriched in the −60 to −25 nt region. CUGCG, a variant of the recurrent CUGMG motif, has also been reported as a YBX1-binding site in the p16INK4a 5 ′ UTR, where YBX1 is associated with increased translation efficiency, particularly under hypoxic conditions (Bisio et al. 2015). These observations suggest that upstream sequence motifs and RNA-binding proteins may contribute to the predictive importance of this region, although the differences in predicted RNA structure were too small to define a clear mechanism.

Module II showed consistent performance on the internal test set and independent proteomic dataset. Its feature contributions differed from those of Module I. Its upstream RNA contribution was concentrated near the AUG and associated predominantly with A/U-rich motifs. Protein sequence contributed most strongly, particularly at the N-terminus, with the largest positional effect within the first 15-20 amino acids. N-terminal residues can influence protein stability through N-degron pathways (Hwang et al. 2010; Timms et al. 2019), providing a possible explanation for this signal. However, peptide detectability in mass spectrometry is also influenced by amino acid composition and physicochemical properties (Zimmer et al. 2018). The N-terminal signal may therefore reflect both protein turnover and sequence features that affect MS detection. Consistent with this complementary role of the two modules, an MS-supported microproteins with literature evidence received a low Module I score but a substantially higher Module II score.

Several limitations should be considered. The labels reflect evidence in currently available datasets rather than the complete translation status of each sORF. Ribo^−^ candidates may be translated in tissues or conditions not represented in public resources, and the background set may contain unrecognized coding sORFs. MS evidence is also influenced by experimental protocols and peptide detectability. Differences between the two tasks may therefore reflect both biological properties and biases in data collection. Moreover, ablation and motif enrichment establish predictive associations, not regulatory mechanisms. Matched Ribo-seq and proteomic datasets, together with targeted mutagenesis, would help distinguish these contributions.

In conclusion, sORF-Trans2MS improves prediction of Ribo-supported sORFs while extending coverage to MS-supported microproteins without public Ribo-seq support. The upstream RNA and N-terminal protein features identified here provide a basis for testing how sequence context influences microprotein translation and detection.

## Material and methods

### Data sources and evidence annotation

Human sORF and microprotein candidates were collected from OpenProt 2.2 (Leblanc et al. 2024), SmProt 2 (Li et al. 2021), Ribotricer (Choudhary et al. 2020), and GENCODE Phase 2 ribosome profiling ORFs (Chothani et al. 2026). Genomic coordinates and transcript annotations were based on the GRCh38 assembly and Ensembl release 106, and the corresponding transcript and genome sequences were obtained from the same Ensembl release (Cunningham et al. 2022). Independent MS-supported microproteins used for external evaluation of Module II were collected from two published proteomic studies (Li et al. 2022a; Li et al. 2026).

Human microprotein annotations were obtained from the GRCh38.106 annotation file of OpenProt 2.2. Only noncanonical sORFs (Altprot) were retained, while reference proteins were excluded. OpenProt entries were grouped according to the MS score and the TE Translation Event score. Entries with MS score equal to 0, 1, and at least 2 were labeled as MS−, MS+, and MS++, respectively. Entries with TE Translation Event score equal to 0, 1 and at least 2 were labeled as Ribo-, Ribo+ and Ribo++, respectively. After transcript matching, sequence extraction, and length filtering, 11,074 Ribo positive OpenProt sORFs encoding 30 to 150 amino acids were retained for Module I. These candidates included 2451 MS^−^Ribo^+^, 1912 MS^−^Ribo^++^, 1787 MS^+^Ribo^+^, 1499 MS^+^Ribo^++^, 1845 MS^++^Ribo^+^, and 1580 MS^++^Ribo^++^ sORFs (**Table 5**). All six evidence groups were included in the Ribo positive candidate pool, regardless of their level of mass spectrometry support.

**Table 5.** Classification of OpenProt sORFs by ribosome profiling and mass spectrometry evidence.

|  | Ribo<sup>-</sup> | Ribo<sup>+</sup> | Ribo<sup>++</sup> |
| --- | --- | --- | --- |
| MS<sup>-</sup> | 182,760 | 2,451 | 1,912 |
| MS<sup>+</sup> | 123,857 | 1,787 | 1,499 |
| MS<sup>++</sup> | 102,930 | 1,845 | 1,580 |

SmProt 2 human sORFs were used only for redundancy filtering during background dataset construction, rather than as training positives, because the exact genome annotation version was unclear and reliable flanking sequence extraction was not feasible (Li et al. 2021).

GENCODE Phase 2 Primary ribosome profiling ORFs were used as an independent dataset for evaluating Module I (Chothani et al. 2026). The original GENCODE Primary dataset contained 10,127 ORFs, of which 6,184 started with ATG. After transcript matching and sequence extraction, 6,178 sORFs were retained for subsequent analysis.

### Construction of the background sORF dataset

Ribotricer was used to obtain candidate sORFs from the GRCh38 index for background dataset construction (Choudhary et al. 2020). To reduce contamination by previously reported microproteins, background sORFs were compared with protein sequences from OpenProt, SmProt, and GENCODE. All sequences were clustered using CD-HIT at 80% amino acid identity (Li and Godzik 2006). Background ORFs that clustered with any known ORF product were removed, while clusters containing only background ORFs were represented by the longest sequence. This filtering retained 759,168 representatives. After transcript matching, flanking sequence extraction, and coordinate deduplication, 75,821 background sORFs were obtained.

### Benchmark dataset construction

For Module I, the 11,074 Ribo positive OpenProt sORFs were used as positive candidates, and the filtered background sORFs were used as negative candidates. To reduce bias related to sequence length, positive and background samples were matched according to ORF length and the available upstream and downstream sequence lengths. Ten matched datasets were generated using different random seeds. Each dataset contained 9,579 Ribo positive sORFs and an equal number of background sORFs. Each dataset was split into training, validation, and test sets at a ratio of 70:15:15 using different random seeds.

For Module II, 102,930 microproteins with strong mass spectrometry evidence but no public Ribo-seq support (MS^++^Ribo^−^) were available as positive candidates. A subset of 15,000 candidates was selected for benchmark construction. These candidates were matched with the background sORFs according to ORF length and the available upstream and downstream sequence lengths. After length matching, 14,697 MS^++^Ribo^−^ candidates were retained and paired with an equal number of background sORFs for Module II training and evaluation. The resulting datasets were also split into training, validation, and test sets at a ratio of 70:15:15 using different random seeds.

### Sequence extraction and preprocessing

For all selected sORFs, upstream, ORF, and downstream sequences were extracted as model inputs. For OpenProt entries, transcript accessions were matched to Ensembl transcript sequences. If one protein accession matched multiple transcripts, the transcript with the longest available sequence was selected. sORF sequences were extracted according to transcript coordinates.

For Ribotricer sORFs, genomic sequences were first extracted according to chromosome coordinates and strand information. sORFs on the negative strand were reverse complemented and then matched back to transcript sequences to obtain 5′-3′ sORF sequences. For GENCODE sORFs, candidate transcripts from the same gene were searched, and the longest transcript containing the ORF sequence was selected.

For each sORF, up to 450 nucleotides upstream of the start codon and up to 450 nucleotides downstream of the stop codon were extracted when available. If the sORF was close to the transcript boundary, only the available sequence was retained. Sequences with missing coordinates, invalid bases, or invalid labels were removed. For the main training benchmarks, only sORFs encoding 30 to 150 amino acids were retained.

### Independent validation dataset of Module I using GENCODE sORFs

GENCODE Phase 2 Primary ribosome profiling sORFs were used to evaluate model generalization. sORF nucleotide and protein sequences were obtained from the GENCODE Phase 2 FASTA files. GENCODE v45 transcript sequences were used for transcript matching and flanking sequence extraction. For each sORF, candidate transcripts from the same gene were searched, and the longest transcript containing the sORF sequence was selected. sORFs without matched transcripts were removed.

After transcript matching and sequence extraction, 6,178 GENCODE ORFs with ATG start were retained as independent candidate sORFs. In addition, 10,000 ribotricer background ORFs that did not appear in any of the ten translation training datasets were sampled as independent controls. The final external validation set contained 16,178 samples. None of the samples in the independent GENCODE test set were used for model training.

### Independent validation dataset of Module II using MS-supported microproteins

An independent mass spectrometry dataset was constructed to evaluate the generalization of Module II. MS-supported microproteins were collected from two published proteomic studies, including 195 candidates from the study by Fengming Li et al.(Li et al. 2026) and 657 candidates from another microproteomics study by Bing Li et al. (Li et al. 2022a). After combining the two datasets, a total of 701 microproteins were retained as independent positive samples. To avoid overlap with the benchmark datasets, sequences overlapping with Module II datasets were removed before external evaluation. Independent negative controls were selected from the filtered Ribotricer background pool and matched to the positive samples according to ORF length and the available upstream and downstream sequence lengths. None of the samples in the independent test set were used for model training.

### RNA-FM and ESM2 feature extraction

RNA sequence representations were extracted using RNA-FM, a pretrained RNA foundation model (Chen et al. 2022). For each sORF, the upstream, ORF, and downstream sequences were encoded separately using the 12-layer RNA-FM model. DNA sequences were converted to RNA by replacing T with U before encoding. Token representations from the final layer were retained, resulting in a 640-dimensional embedding for each nucleotide. Special tokens introduced by the model were excluded from the saved representations. The ORF sequence included the terminal stop codon during RNA encoding.

Protein sequence representations were extracted using ESM-2 (Lin et al. 2023). ORF nucleotide sequences were translated into amino acid sequences after removal of the terminal stop codon. The esm2_t33_650M_UR50D model was used, and residue representations from the final layer were retained. Special tokens at both ends of the sequence were removed, resulting in a 1280-dimensional embedding for each amino acid residue.

### Model design of sORF-Trans2MS

sORF-Trans2MS was developed as a dual-module deep learning framework for Ribo-supported sORF translation and MS-supported microprotein prediction (**Figure 1)**. **Figure 1A** summarizes the construction of the Ribo^+^ and MS^++^Ribo^−^ benchmark datasets together with the filtered background sORF set. Module I uses RNA-FM token representations from the upstream, ORF, and downstream regions together with relative position information, followed by a Transformer encoder and an MLP classifier for Ribo-supported translation prediction (**Figure 1B**). Module II integrates RNA-FM representations of the three RNA regions with ESM-2 residue representations of the encoded protein. These features are processed by separate multi-scale CNN encoders, projected into RNA and protein representations, concatenated, and passed to an MLP classifier for MS-supported microprotein prediction (**Figure 1C**). The two modules were trained independently and generated separate prediction probabilities for their corresponding tasks.

### Module I model construction

Module I was designed to predict Ribo-supported sORF translation from the upstream, ORF, and downstream RNA regions (**Figure 1B**). For each nucleotide, the RNA-FM representation with 640 dimensions was combined with four relative position features, including the normalized position, squared position, and sine and cosine transformations of the position. The resulting features were projected into a 256-dimensional hidden space using a linear layer followed by layer normalization, GELU activation, and dropout. Region embeddings were then added to distinguish tokens from the upstream, ORF, and downstream regions. The three regions were concatenated and jointly processed by a two-layer Transformer encoder with 8 attention heads and a feedforward dimension of 512. Padding positions were excluded using attention masks. The encoded tokens were summarized by masked mean and max pooling, and the two pooled vectors were concatenated into a 512-dimensional representation. This representation was passed through an MLP with dimensions of 512, 256, 128, and 1 to generate the final prediction score. Dropout was set to 0.2.

### Module II model construction

Module II was designed to predict MS^++^Ribo^−^ microproteins by integrating RNA context with protein sequence information (**Figure 1C**). RNA-FM representations from the upstream, ORF, and downstream regions and ESM-2 residue representations of the encoded protein were processed by separate CNN encoders. Four relative position features were appended to each token representation before encoding. Each encoder first projected the input features into a 128-dimensional hidden space and then applied parallel one-dimensional convolutional branches with kernel sizes of 3, 5, and 7. Residual convolutional blocks with layer normalization were used within each branch, and mean and max pooling were applied to summarize the encoded sequence. The three convolutional branches produced a combined 768-dimensional representation for each RNA region and protein sequence. The three RNA representations were concatenated and projected into a 256-dimensional RNA representation, while the protein representation was independently projected to 256 dimensions. The resulting RNA and protein representations were concatenated into a 512-dimensional vector and passed through an MLP with dimensions of 512, 256, 128, and 1 to generate the final prediction score. Dropout was set to 0.3.

### Model training and implementation

Module I was implemented in Python using PyTorch and trained as a binary classifier. Model parameters were optimized using AdamW with an initial learning rate of 1*10^-4^ and a weight decay of 5*10^-4^. The model was trained with a batch size of 64 for a maximum of 50 epochs. Binary cross entropy loss with class weighting based on the ratio of negative to positive samples was used for optimization. A ReduceLROnPlateau scheduler reduced the learning rate by a factor of 0.5 when validation AUROC (Area Under the Receiver Operating Characteristic Curve) did not improve for three consecutive epochs. Early stopping was applied with a patience of 10 epochs, and the model with the highest validation AUROC was retained for test-set evaluation. Gradient clipping with a maximum norm of 1.0 was applied during training. Ten independent runs were performed using random seeds 42-51, and internal test performance was summarized as the mean ± standard deviation across the ten runs.

Module II was also implemented in Python using PyTorch and trained as a binary classifier. AdamW was used with an initial learning rate of 1*10^-4^ and a weight decay of 5*10^-4^. The model was trained with a batch size of 512 for a maximum of 50 epochs using binary cross-entropy loss. A ReduceLROnPlateau scheduler monitored validation AUROC and reduced the learning rate by a factor of 0.5 after three epochs without improvement, with a minimum learning rate of 1*10^-6^. Early stopping was applied with a patience of 10 epochs, and the checkpoint with the highest validation AUROC was selected for test-set evaluation. Gradient clipping with a maximum norm of 1.0 was used during optimization. Ten independent runs were performed using random seeds 42–51, and internal test performance was summarized as the mean ± standard deviation across the ten runs. All experiments were run on a Linux server equipped with an NVIDIA GeForce RTX 3090 GPU (24 GB memory).

### Evaluation metrics

Model performance was evaluated using accuracy, precision, recall, F1 score, area under the receiver operating characteristic curve (AUROC), and area under the precision–recall curve (AUPR). Accuracy, precision, recall, and F1 score were calculated from binary predictions using a probability threshold of 0.5, whereas AUROC and AUPR were calculated from continuous prediction scores. Let TP, TN, FP, and FN denote true positives, true negatives, false positives, and false negatives, respectively.

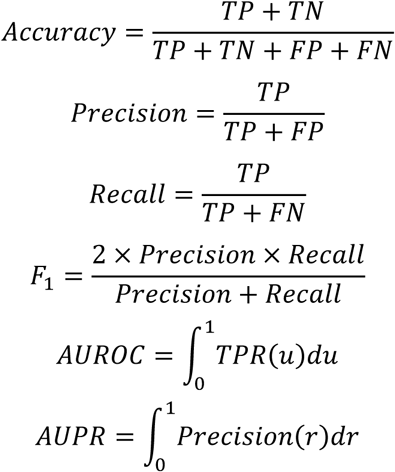

AUROC measures the ability of a model to rank positive and negative samples across classification thresholds, whereas AUPR summarizes the precision-recall relationship and emphasizes positive-class retrieval. For the internal benchmark datasets, performance was summarized as the mean ± standard deviation across ten independent runs.

### Architecture ablation analysis

Architecture ablation analysis was performed for Module I using the first five independently generated matched datasets. For each dataset, we evaluated the full model and three ablated variants in which the relative position features, region embeddings, or Transformer encoder were removed, respectively. All other training settings were kept unchanged. Performance was summarized as the mean across the five runs.

### RNA feature ablation analysis

RNA feature ablation was performed separately for Module I and Module II using the test sets from all ten runs. For region level analysis, RNA-FM representations of the upstream, ORF, or downstream region were individually set to zero while the position features were retained. In Module II, the protein branch remained unchanged during RNA ablation. For position level analysis, a 9 nt sliding window with a stride of 3 nt was applied across the three RNA regions, and the RNA-FM representations within each window were set to zero. Feature importance was quantified by the decrease in AUROC and AUPR after ablation:

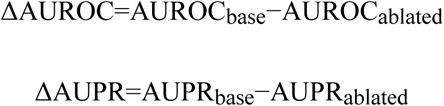

Larger positive values indicate a greater contribution to model performance. Results were summarized across ten runs. Paired comparisons between ablation conditions were performed using two-sided Wilcoxon signed-rank tests, followed by Benjamini–Hochberg correction for multiple testing.

### Protein feature ablation analysis

Protein feature ablation was performed on Module II using the fixed test sets from all ten runs. During protein ablation, the RNA branch was kept unchanged. Ablation was conducted at three levels: the entire protein sequence, the N-terminal and C-terminal halves, and local sequence windows.

For half-sequence analysis, the protein representation was divided into N-terminal and C-terminal halves and each half was independently set to zero. For position level analysis, a 5 amino acid sliding window with a stride of 1 was applied separately within the first 75 positions from the N terminus and the last 75 positions from the C terminus. Protein importance was quantified using ΔAUROC and ΔAUPR as defined above. Results were summarized across ten runs, and paired comparisons were evaluated using two-sided Wilcoxon signed-rank tests with Benjamini Hochberg correction.

### Upstream RNA motif enrichment analysis

Upstream sequences were divided into five regions relative to the AUG start codon: −120 to - 85, −96 to −61, −60 to −25, −36 to −1, and −24 to −1 nt. Differential motif enrichment between positive and negative samples was analyzed using STREME from the MEME Suite (Bailey et al. 2015; Bailey 2021). The analysis was performed with the RNA alphabet, motif widths of 5–10 nt, and a second-order background model. Motifs with E-values ≤ 0.05 were considered significant. De novo motifs were identified from run 00 and visualized using sequence logos generated by the MEME Suite. For selected Module I motifs, enrichment was additionally assessed across ten independently sampled datasets using odds ratios and 95% confidence intervals.

## Data and Code Availability

The code and processed datasets are available at https://github.com/stevencapeng/sORF_Trans2MS. Supplemental material is available for this article.

## Acknowledgements

This project was supported by the National Key Research and Development Program of China (no. 2021YFA1302603), the Shenzhen Science and Technology Program (no. QTD20221101093558015), the Shenzhen Key Laboratory of Functional Proteomics (no. ZDSYS20230626090803004), and the Guangdong Engineering Technology Research Center of Proteomic Analysis and Sequencing (no. 2023B190). Schematic diagrams are created with BioRender.com. The research work in Hu’s lab was supported by the National Natural Science Foundation of China (82173131 to H.H.); and the Natural Science Foundation of Guangdong Province (2023A1515010124 and 2024A1515012783 to H.H.)

## Author Contributions

J.H. conceived and developed the computational framework, performed the analyses, prepared the figures, and drafted the manuscript. J.S. contributed to data collection and preparation. Q.C. contributed to dataset preparation and model optimization. H.H. contributed to study supervision and manuscript revision. C.S.H.T. conceived and supervised the study and revised the manuscript. All authors read and approved the final manuscript.

