## Supplemental material for "sORF-Trans2MS: a two-module deep learning framework for sORF translation and MS-supported microprotein prediction"

### Supplementary material

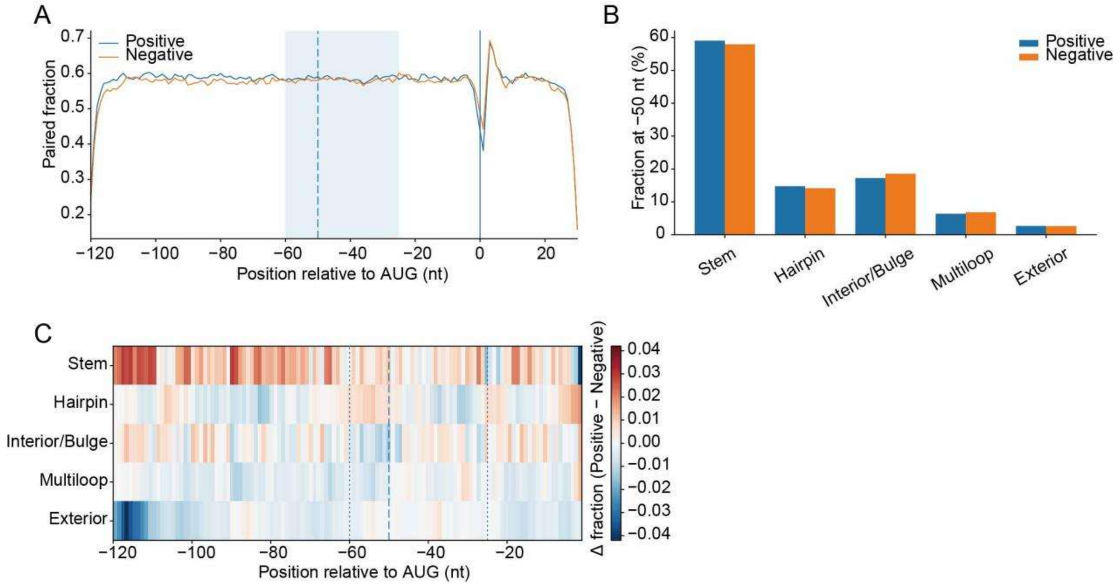


**Figure S1. Comparison of predicted RNA secondary structures between positive and negative sORFs in Module I dataset.**

**A**. Position-wise paired fraction from RNAfold minimum free energy structures. The shaded region marks -60 to -25nt, the dashed line marks -50nt, the solid line indicates the AUG start site. **B**. Bar plot shows the fractions of different secondary structure at -50nt. **C**. Position-wise differences in frequency of each RNA structure between positive and negative sORFs (positive minus negative).

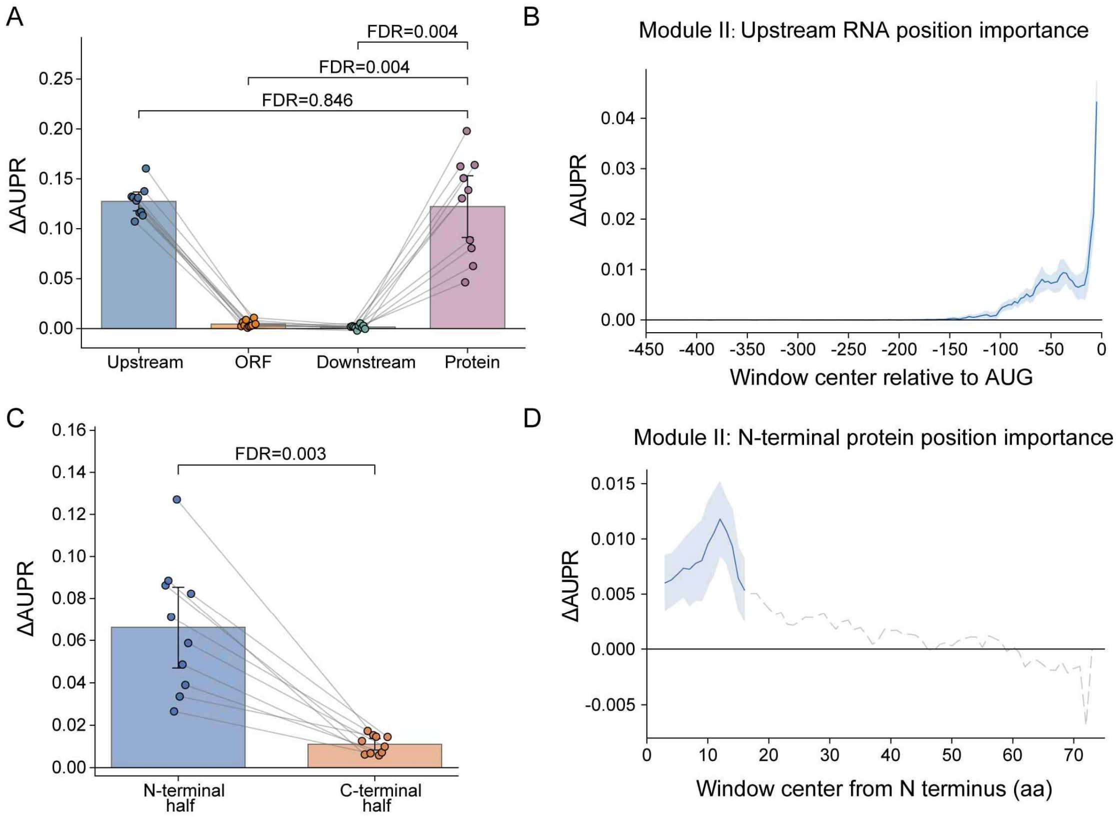


**Figure S2. RNA and protein ablation analysis of Module II.**

**A**. Comparison of ΔAUPR after ablation of the upstream, ORF, downstream, and protein inputs. **B**. Position-level ΔAUPR across the upstream RNA region. **C**. Comparison of ΔAUPR after ablation of the N-terminal and C-terminal halves of the protein sequence. **D**. Position-level ΔAUPR across the N-terminal half of the protein sequence. All of the results are summarized across ten runs. Positive ΔAUPR values indicate decreased model performance after ablation and therefore greater predictive importance. Statistical comparisons were performed using paired two-sided Wilcoxon signed-rank tests with Benjamini-Hochberg correction.

**Table S1. Enriched upstream sequence motifs identified in Module I.**

| Region | Rank | Motif | E-value | Sites |
|---|---|---|---|---|
| −120 to −85 | 1 | SGCSGCGSS | 2.2e-16 | 1973 |
| −120 to −85 | 2 | VSCGG | 3.3e-12 | 3999 |
| −120 to −85 | 3 | CGGAA | 0.18 | 136 |
| −120 to −85 | 4 | GCGCC | 2 | 43 |
| −120 to −85 | 5 | GRKCG | 3 | 45 |
| −120 to −85 | 6 | AACCCACAA | 5.2 | 27 |
| −96 to −61 | 1 | CGCGSCSG | 7.7e-17 | 2008 |
| −96 to −61 | 2 | CGCCG | 0.0026 | 3508 |
| −96 to −61 | 3 | CGGDS | 0.051 | 231 |
| −96 to −61 | 4 | CGGAA | 0.11 | 162 |
| −96 to −61 | 5 | UCAGCWGC | 1.2 | 145 |
| −96 to −61 | 6 | CGUUAAA | 1.7 | 29 |
| −96 to −61 | 7 | CUGAGUUG | 1.7 | 45 |
| −60 to −25 | 1 | CCGCCGCSG | 6.4e-15 | 1515 |
| −60 to −25 | 2 | CCGGH | 4.8e-06 | 3580 |
| −60 to −25 | 3 | CUGMG | 0.046 | 961 |
| −60 to −25 | 4 | CUGAA | 0.88 | 817 |
| −60 to −25 | 5 | AGCUAGAA | 3.7 | 65 |
| −60 to −25 | 6 | CGYUG | 4.7 | 268 |
| −36 to −1 | 1 | CCGCCGCSG | 4e-13 | 1328 |
| −36 to −1 | 2 | CCUGA | 5e-06 | 4079 |
| −36 to −1 | 3 | GCGCG | 0.016 | 1821 |
| −36 to −1 | 4 | CGUUUC | 1.4 | 163 |
| −36 to −1 | 5 | AGAUAACC | 3 | 40 |
| −36 to −1 | 6 | AKAAGGWAG | 4.5 | 55 |
| −24 to −1 | 1 | SCGCG | 2.4e-08 | 2908 |

| | | | | |
|---|---|---|---|---|
| −24 to −1 | 2 | CCURA | 2.7e-06 | 2104 |
| −24 to −1 | 3 | UAGGAR | 0.21 | 480 |
| −24 to −1 | 4 | SAGURA | 0.5 | 283 |
| −24 to −1 | 5 | GGUAAC | 1.2 | 98 |
| −24 to −1 | 6 | ACCGUACA | 1.5 | 23 |

**Table S2. Enriched upstream sequence motifs identified in Module II.**

| Region | Rank | Motif | E-value | Sites |
|---|---|---|---|---|
| −120 to −85 | 1 | RBUUAG | 0.056 | 1306 |
| −120 to −85 | 2 | GAAGCGU | 0.062 | 38 |
| −120 to −85 | 3 | UACUAA | 0.069 | 804 |
| −120 to −85 | 4 | UWAAG | 0.23 | 6474 |
| −120 to −85 | 5 | AAAUAUUA | 0.25 | 2015 |
| −120 to −85 | 6 | UUUUCUU | 5 | 890 |
| −120 to −85 | 7 | GCAGCUCUGA | 5.5 | 39 |
| −120 to −85 | 8 | AAGCGACU | 8 | 52 |
| −96 to −61 | 1 | UARHHH | 0.0043 | 7400 |
| −96 to −61 | 2 | AUUCUU | 0.15 | 172 |
| −96 to −61 | 3 | UUUUGA | 0.26 | 4286 |
| −96 to −61 | 4 | UGGAAACCAA | 1.1 | 31 |
| −96 to −61 | 5 | CCUCUGCAA | 1.9 | 28 |
| −96 to −61 | 6 | CCCAGCUAU | 3 | 42 |
| −60 to −25 | 1 | WWHUAR | 2.4e-11 | 7272 |
| −60 to −25 | 2 | NHBUGA | 1.3e-05 | 4926 |
| −60 to −25 | 3 | AAGGACAGU | 0.07 | 75 |
| −60 to −25 | 4 | AAUUU | 0.27 | 2223 |
| −60 to −25 | 5 | AAAAG | 0.39 | 1802 |
| −60 to −25 | 6 | GGCYGAGGC | 0.41 | 86 |

| | | | | |
|---|---|---|---|---|
| −60 to −25 | 7 | UARDH | 1.9 | 665 |
| −60 to −25 | 8 | AACCCCAUCU | 4.5 | 59 |
| −60 to −25 | 9 | WCCAGCCUGG | 4.5 | 45 |
| −36 to −1 | 1 | UAAAW | 6e-15 | 10611 |
| −36 to −1 | 2 | HDUAR | 3.3e-10 | 795 |
| −36 to −1 | 3 | DKUGA | 0.0084 | 1622 |
| −36 to −1 | 4 | KCAUGAGC | 0.054 | 49 |
| −36 to −1 | 5 | UGUCUCAAAA | 5.7 | 35 |
| −36 to −1 | 6 | CAGCUUUACA | 6.4 | 88 |
| −36 to −1 | 7 | AACAAAUACA | 7 | 33 |
| −24 to −1 | 1 | HDUAA | 1.7e-18 | 8622 |
| −24 to −1 | 2 | UGAAA | 0.0054 | 1318 |
| −24 to −1 | 3 | GGCUGA | 0.19 | 83 |
| −24 to −1 | 4 | CAAAUUGCA | 1.1 | 34 |
| −24 to −1 | 5 | UGGAGGUUGC | 3 | 22 |
| −24 to −1 | 6 | UAAGR | 3.7 | 116 |
